# Universal metabolic constraints on the thermal tolerance of marine phytoplankton

**DOI:** 10.1101/358002

**Authors:** Samuel Barton, James Jenkins, Angus Buckling, C.-Elisa Schaum, Nicholas Smirnoff, Gabriel Yvon-Durocher

## Abstract

Marine phytoplankton are responsible for over 45% of annual global net primary production. Ocean warming is expected to drive massive reorganisation of phytoplankton communities, resulting in pole-ward range shifts and sharp declines in species diversity, particularly in the tropics. The impacts of warming on phytoplankton species depend critically on their physiological sensitivity to temperature change, characterised by thermal tolerance curves. Local extinctions arise when temperatures exceed species’ thermal tolerance limits. The mechanisms that determine the characteristics of thermal tolerance curves (e.g. optimal and maximal temperatures) and their variability among the broad physiological diversity of marine phytoplankton are however poorly understood. Here we show that differences in the temperature responses of photosynthesis and respiration establish physiological trade-offs that constrain the thermal tolerance of 18 species of marine phytoplankton, spanning cyanobacteria as well as the red and green super-families. Across all species we found that rates of respiration were more sensitive to increasing temperature and typically had higher optimal temperatures than photosynthesis. Consequently, the fraction of photosynthetic energy available for allocation to growth (carbon-use efficiency) declined exponentially with rising temperatures with a sensitivity that was invariant among the 18 species. Furthermore, the optimal temperature of growth was generally lower than that of photosynthesis and as a result, supra-optimal declines in growth rate were associated with temperature ranges where the carbon-use efficiency exhibited accelerated declines. These highly conserved patterns demonstrate that the limits of thermal tolerance in marine phytoplankton are underpinned by common metabolic constraints linked to the differential temperature responses of photosynthesis and respiration.

**Significance Statement:** The impacts of warming on marine phytoplankton depend on their sensitivity to rising temperatures, yet there is currently limited understanding of the mechanisms that limit thermal tolerance among the diversity of marine phytoplankton. Using a comparative study on the dominant, ecologically important lineages of marine phytoplankton – Bacillariophyceae, Dinophyceae, Cyanophyceae, Prasinophyceae, Prymnesiophyceae – we show that rates of respiration are consistently more sensitive to increasing temperature than photosynthesis. Consequently, the fraction of photosynthetic energy available for growth declines with rising temperatures with a sensitivity that is invariant among species. Our results suggest that declines in phytoplankton performance at high temperatures are driven by universal metabolic constrains linked to rising respiratory costs eventually exceeding the supply of reduced carbon from photosynthesis.

## Introduction

The planet’s oceans are changing at an unprecedented rate (1); over the past half-century average sea surface temperatures have been increasing by 0.1 °C per decade (2) and are projected to rise by a further 3°C or more by the end of the century (3). Ocean warming is thought to be a key driver of recent declines in phytoplankton productivity (4–6), and models of marine biogeochemistry predict further reductions in productivity over the 21st century as temperatures exceed limits of thermal tolerance and nutrient limitation increases in warmer, more stratified oceans (7). Thermal tolerance curves of marine phytoplankton (like all ectotherms) exhibit characteristic unimodality and left-skew, meaning that fitness declines more sharply above the optimum temperature than below (8). Marine phytoplankton species exhibit substantial variability in their thermal tolerance. Optimal temperatures for growth range between approximately 2 to 38°C and are positively correlated with the average temperature of the environment, indicating a global pattern of thermal adaptation (8, 9). Ocean warming is expected to result in major reorganisation of marine phytoplankton communities as temperatures exceed the thermal optima of some species but not others. In particular, tropical and sub-tropical regions are projected to experience pronounced declines in species diversity and productivity (8, 9) because many of the taxa in these areas already exist close to their limits of thermal tolerance. Despite its importance for predicting the impacts of global warming on marine phytoplankton communities, we currently understand very little about the physiological processes that determine the limits of thermal tolerance in marine phytoplankton.

To address this fundamental knowledge gap we carried out a large-scale experiment to investigate the physiological mechanisms that set the limits of thermal tolerance in marine phytoplankton. Our experiments span a representative sample of the broad physiological and phylogenetic diversity of the marine phytoplankton; including 18 species belonging to ecologically important functional groups – Cyanobacteria, Diatoms, Dinoflagellates, Coccolithophores, Rhodophytes, Chlorophytes and Prasinophytes (Table 1, SI). These species were chosen to encompass the putative primary and secondary endosymbionts of both the red and green super-families, and thus reflect the complex evolutionary histories of marine phytoplankton (10, 11). This allowed for us to investigate whether, in spite of such physiological diversity in plastid evolution, similar physiological constraints underpin the limits of thermal tolerance.

## Results & Discussion

We first characterised variability in thermal tolerance curves among taxa by measuring growth rates for each species across a temperature gradient spanning 15 to 37°C and fitting the Sharpe-Schoolfield equation for high temperature inactivation to the data using non-linear mixed effects modelling (12, 13) (Fig. 1). The upper limits of thermal tolerance varied across the taxa, with *T_max_^μ^* (maximum temperature of observed growth), ranging from 27°C to 37°C. The optimal temperature of growth, *T_opt_^μ^*, ranged from 23.8°C to 34.0°C and the activation energy, *E_a_^μ^* – which characterises the increase in rate up to *T_opt_^μ^* – ranged from 0.40 eV to 1.46 eV, with an average *E_a_^μ^* of 0.77eV (95% CI: 0.58 to 0.97) (Fig. 1, and Table 2, SI). These *E_a_^μ^* values highlight that the temperature dependence of growth at the species level is significantly higher than previously reported temperature dependence parameters, such as the canonical Eppley coefficient (equivalent to *E_a_^μ^* ≈ 0.3 eV), that are derived by comparing maximum growth rates across many species and are the standard way in which the impacts of warming on phytoplankton productivity are represented in models of marine biogeochemistry (14–16). These findings suggest that the Eppley coefficient (and other values from similar analyses (16)), which capture the broad-scale, macroecological impacts of temperature along geographic gradients, might significantly under estimate the impacts of temperature fluctuations on phytoplankton growth at local to regional scales (see Supplementary Information: additional text, S1, for further discussion).

**Fig. 1.**
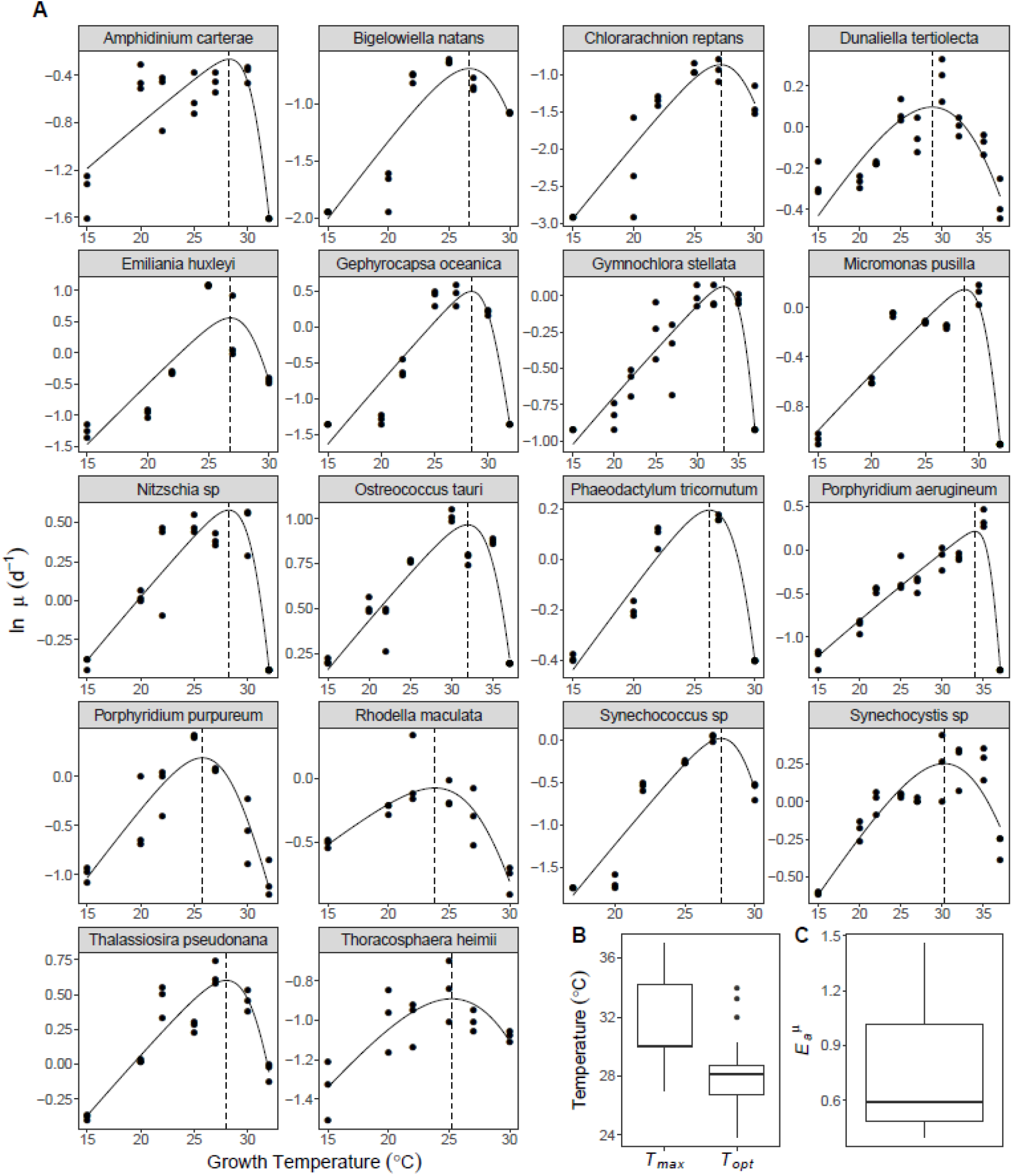
Thermal tolerance curves for 18 species of marine phytoplankton. **(A)** Thermal reaction norms for all 18 species used in this study. The data points presented are the natural logarithm of per capita growth rate, *μ*, for each replicate (n= minimum of 3 technical replicates per assay temperature for each species). The fitted lines are from the predicted random effect of species derived from non-linear mixed effects modelling with a Sharpe-Schoolfield model. The vertical dashed lines correspond with the optimal temperatures of growth. **(B)** Boxplot distributions of optimal growth temperatures (*T_opt_*) and maximum temperatures of growth (*T_max_*) across all 18 species. **(C)** Boxplot distribution of growth activation energy, or temperature dependence (*E_a_^μ^*), across all 18 species (Table 2, SI). The bold horizontal line corresponds to the median value, the top and bottom of the box correspond to the 75^th^ and 25^th^ percentiles and the whiskers extend to the largest and smallest values no greater or less than 1.5 × the interquartile range, beyond which the points are plotted as outliers.

To determine the physiological processes that shape the thermal tolerance curves, in particular those that determine the optimum temperature and supra-optimal declines in growth rate, it is essential to understand how the key metabolic pathways that drive biomass synthesis respond to warming. Despite having diverse evolutionary histories, all unicellular phytoplankton share common, key metabolic pathways (17) and their ability to sequester carbon, and therefore grow, is ultimately determined by photosynthesis and respiration (18, 19). The light-dependent reactions of photosynthesis account both for the processes that convert inorganic carbon to organic carbon stores and those that facilitate the production of ATP and reductant used to fuel biomass synthesis (20). The dark reactions in respiration can be conceptually divided into ‘growth’ and ‘maintenance’ components (18–21). ‘Growth-respiration’ provides the ATP, reductant and carbon skeletons required for producing new biomass and is expected to be proportional to the rate of growth. By contrast, ‘maintenance-respiration’ provides the ATP for macromolecular turnover and the maintenance of solute gradients, and is proportional to cell biomass (20). Whilst dark respiration clearly plays an important role in photolithotrophic growth in microalgae, the majority of the energy used to fuel biosynthesis (between 60 – 90%) is thought to derive from photosynthesis (20, 21). To understand the physiological constraints that shape the variability in phytoplankton thermal tolerance, we quantified temperature-dependent variation in rates of photosynthesis and dark respiration in the 18 species of marine phytoplankton.

For each species, we measured the acute responses of gross photosynthesis and dark respiration across a temperature gradient spanning 7°C to 49°C, and quantified the resultant thermal response curves by fitting the Sharpe-Schoolfield equation for high temperature inactivation to the data using non-linear mixed effects modelling (see Methods). We found consistent differences in the parameters characterising the thermal responses of photosynthesis and respiration across all the species in this study despite their diverse evolutionary histories (Fig. 2, Fig. 3 and 4, SI). The activation energy for respiration was greater than that of photosynthesis (i.e. *E_a_^R^ > E_a_^p^*; Fig. 3B, Fig. 3 and 4, SI) in all 18 species. Pooling the parameters across species yielded an average activation energy for photosynthesis of *E_a_^p^*= 0.74 eV (95% CI: 0.69 to 0.79), whilst the average for respiration was *E_a_^R^*= 1.07 eV (95% CI: 0.98 to 1.15). Critically, the average activation energy for photosynthesis was statistically indistinguishable from that of growth rate (*E_a_^μ^* = 0.77eV, 95% CI: 0.58 to 0.97). These results demonstrate that respiratory costs become an increasingly large proportion of photosynthetic carbon fixation and biomass synthesis as temperatures rise toward the peak of the thermal response curves. We also found that for most species, the optimum temperature for respiration was higher than that of photosynthesis (i.e. *T_opt_^R^ > T_opt_^p^*), with the average thermal optimum for photosynthesis, *T_opt_^p^* = 31.18°C ± 0.83 (s.e.m.) and respiration, *T_opt_^R^* = 32.91°C ± 0.48 (s.e.m.) (Fig. 3C, Fig. 3 and 4, SI). Furthermore, in all species, the deactivation energy, which characterises the speed that rates decline past the optimum, was lower for respiration relative to photosynthesis (i.e. *E_h_^p^ > E_h_^R^*), with the average across species for photosynthesis *E_h_^p^* = 6.08 (95% CI: 5.04 to 7.12) and respiration *E_h_^R^* = 2.62 (95% CI: 2.31 to 2.93) (Fig. 3D, Fig. 3 and 4, SI). Thus, as temperatures rise beyond *T_opt_*, rates of photosynthesis decline faster than rates of respiration. Overall these findings show remarkable consistency across diverse taxa (Fig. 3 and 4, SI) in how differences in the parameters that characterise the thermal responses of photosynthesis and respiration result in increasing respiratory expenditure of carbon fixed by photosynthesis as temperatures rise.

**Fig. 2.**
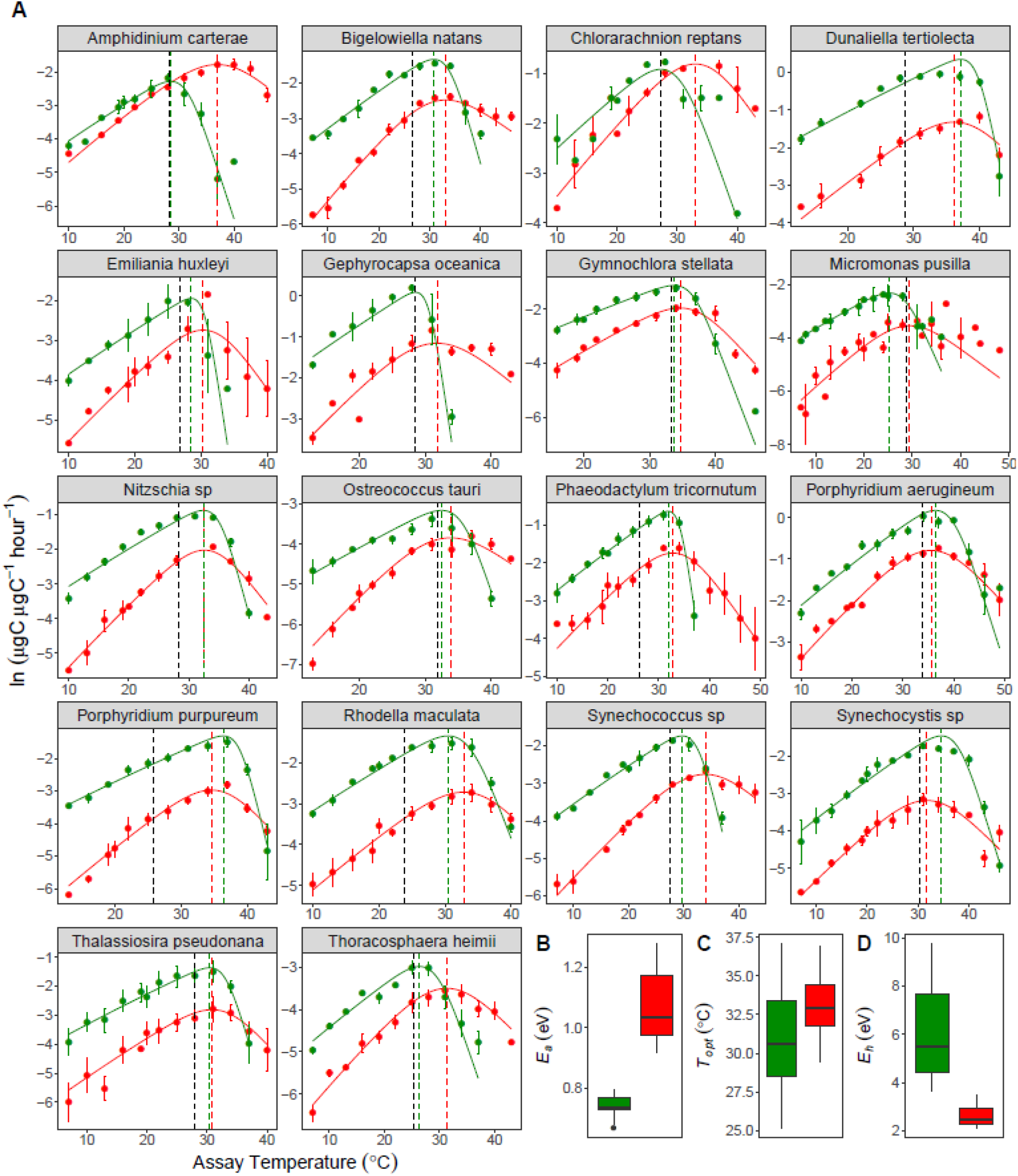
Thermal performance curves for respiration and gross photosynthesis in 18 species of marine phytoplankton. **(A)** Metabolic thermal performance curves for all 18 species used in this study. Green colouring denotes gross photosynthesis, red colouring denotes respiration. The data points presented are the natural logarithm of mean metabolic, with error bars denoting ± s.e.m (n =minimum of 3 biological replicates per response for each species). The fitted lines for each species are from the random effects of a non-linear mixed effects model fitted to the rate data using the Sharpe-Schoolfield equation (see Methods). The vertical dashed lines correspond with the optimal temperatures for each metabolic flux, with the black dashed line added to show optimal growth temperature. (**B, C and D**) Boxplots showing the distribution of the estimated values for activation energy (*E_a_*), optimal temperature (*T_opt_*) and deactivation energy (*E_h_*) for photosynthesis and respiration across the 18 species (Tables 4 and 5, SI). The bold horizontal line corresponds to the median value, the top and bottom of the box correspond to the 75^th^ and 25^th^ percentiles and the whiskers extend to the largest and smallest values no greater or less than 1.5 × the interquartile range, beyond which the points are plotted as outliers.

**Figure. 3.**
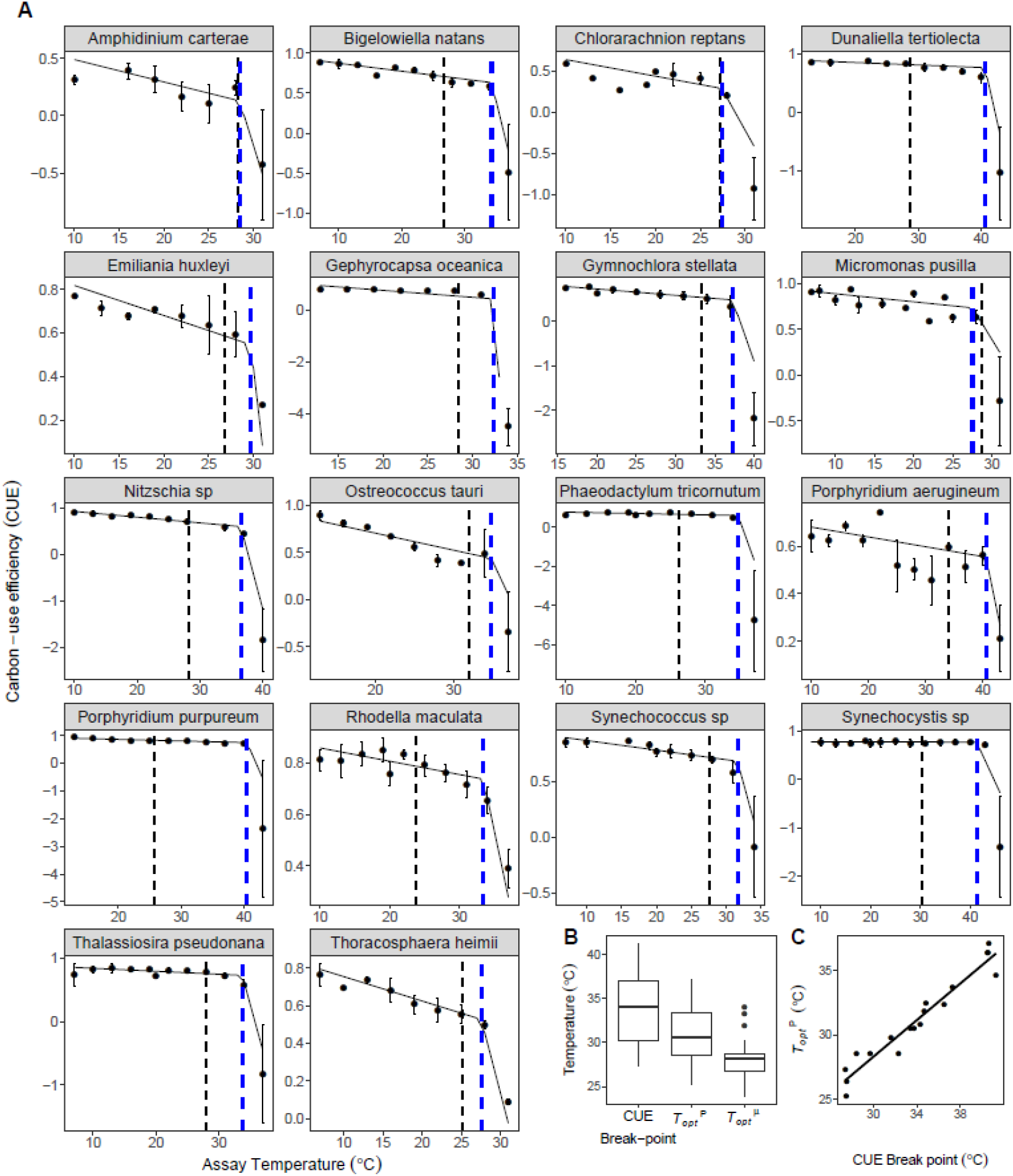
Carbon-use efficiency breakpoints constrain the optimal temperature of growth. **(A)** Segmented linear regression models fitted to the predicted carbon use efficiency (CUE), derived from the thermal performance parameters of respiration and photosynthesis for each species (Fig 2). The modelled response is presented here alongside the calculated mean CUE at each assay temperature, with with error bars denoting ± s.e.m (n =minimum of 3 biological replicates per response for each species). The dashed vertical dashed blue line represents the predicted break-point in the model, where there was a significant change in the slope of the CUE thermal response. The dashed vertical black line represents the estimate optimal temperature of growth (Fig. 1). In most cases this either coincides with the break-point, falling within the 95% CIs of the break-point, or was lower than the break-point. **(B)** Boxplots showing the distribution of the estimated values for the CUE break-point temperature, optimal temperature of gross photosynthesis (*T_opt_^p^*) and optimal temperature of growth (*T_opt_^μ^*) across the 18 species (Tables 2, 4 and 5, SI). The bold horizontal line corresponds to the median value, the top and bottom of the box correspond to the 75^th^ and 25^th^ percentiles and the whiskers extend to the largest and smallest values no greater or less than 1.5 × the interquartile range, beyond which the points are plotted as outliers. **(C)** The significant coupling between the CUE and *T_opt_^p^*, illustrating that the sharp declines in CUE are determined by the universal metabolic constrains identified in Fig. 2.

The carbon-use efficiency (CUE = *1-R/P*), is an estimate of the fraction of photosynthetic energy (P) that can be allocated to growth after accounting for respiration (R). Recent work on both marine and freshwater phytoplankton species suggests that declines in CUE at high temperature may be linked to impaired performance at supra-optimal temperature (22, 23). Furthermore, observations that the evolution of elevated thermal tolerance are coupled with adaptive shifts in metabolic traits that increase CUE at high temperature (22–24), imply an important role for CUE in constraining thermal tolerance that could provide a general explanation for high-temperature impairment of growth across the diversity of the phytoplankton. To determine whether the differential thermal responses of photosynthesis and respiration can help explain the physiological processes that constrain the thermal tolerance curves of diverse phytoplankton, we quantified how the CUE varied as a function of temperature. Consistent with previous work, we found that the CUE decreased with increasing temperature in all 18 species. Declines in the CUE with rising temperature were however highly non-linear, with the fall in CUE dramatically accelerating at high temperatures. Because *T_opt_^R^ > T_opt_^p^* and *E_h_^p^ > E_h_^R^* for most species, as temperature rose beyond *T_opt_^p^* the CUE exhibited an accelerated decline at high temperatures. To quantify this non-linear response and the location of the inflection point where declines in CUE become accelerated, we fitted a break-point model to the thermal responses of the CUE. We found a significant break-point in the thermal response of the CUE for all 18 species that was tightly coupled with *T_opt_^p^* (Fig. 3). As *E_a_^R^ > E_a_^p^* for all species, temperature dependent declines in CUE up to the break-point were universal across the species (Fig. 4) with an average activation energy, *E*_a_^CUE^, of -0.12eV (95% CI: -0.16 to -0.08). Furthermore, in all 18 species the optimum temperature for growth *(T*_opt_^μ^) either coincided with the CUE break-point (i.e. the 95% CIs of the CUE break-point included *T_ċpt_^μ^*), or was lower than the CUE break-point (Fig. 3). This finding suggests that temperature-driven declines in the CUE, linked to fundamental differences in the intrinsic thermal responses of photosynthesis and respiration, could play an important role in constraining the thermal tolerance of diverse marine phytoplankton. Because the metabolic costs for repair and maintenance are largely accounted for by dark respiration (20, 21) the temperature-driven declines in the CUE likely reflect increases in the costs associated with maintenance and repair of heat-induced cellular damage that eventually exceed the rate of substrate supply by photosynthesis, causing rates of growth to decline at supra-optimal temperatures.

**Figure. 4.**
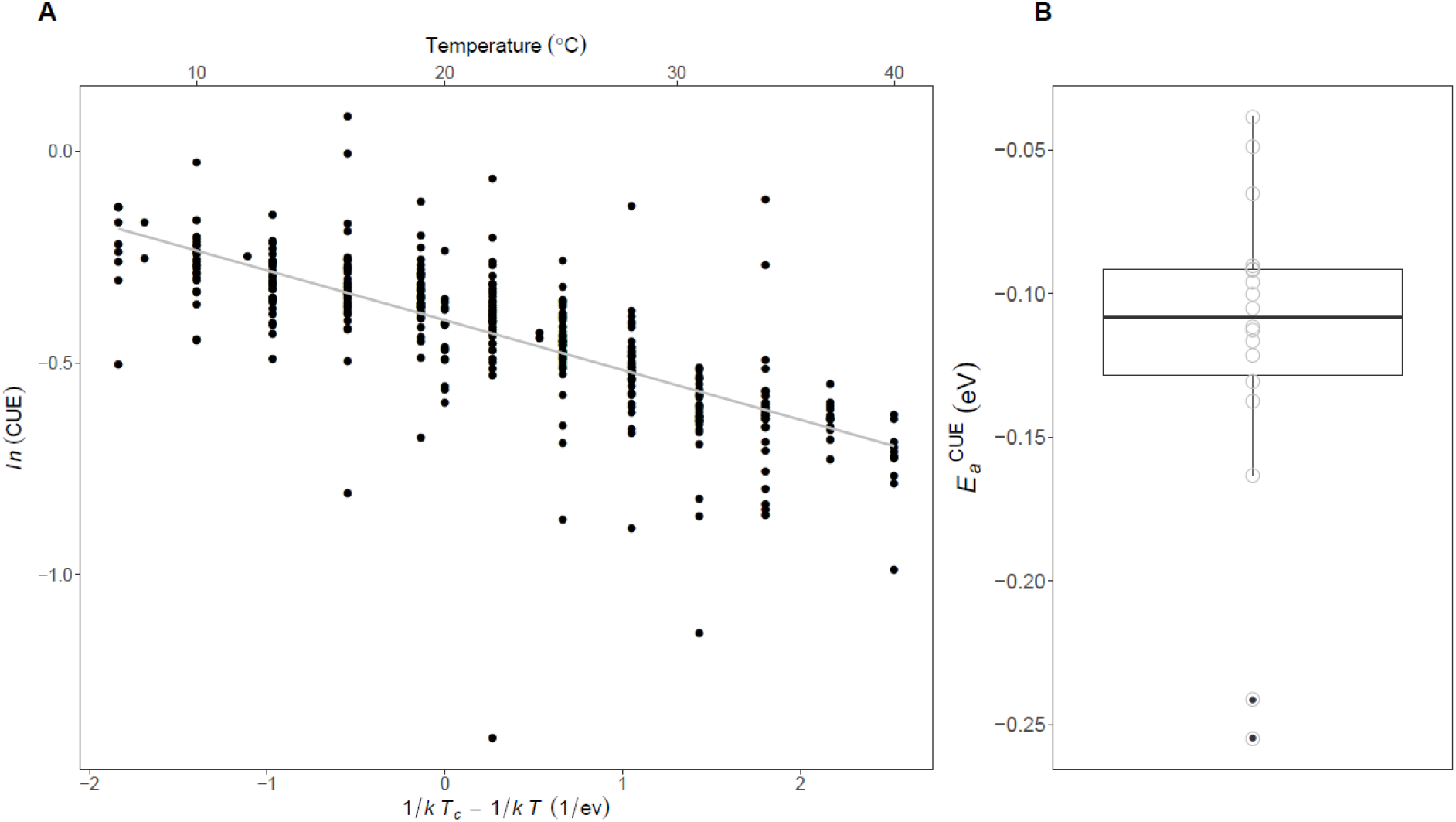
The temperature dependence of the carbon-use efficiency. **(A)** A scatterplot showing the relationship between the natural logarithm of the carbon-use efficiency (CUE) and standardised Boltzmann temperature up to the CUE break-point (Fig. 3) for the pooled dataset of 18 species, where *T_c_* = 20°C and *k* is the Boltzmann constant (8.62 × 10^−5^ eV). The fitted line represents the fixed effect of a linear mixed effects model fitted to the data using the Boltzmann-Arrhenius equation (see Methods). Values of ln(CUE) have been standardised by dividing by the species-specific intercept derived from the random effects of the mixed effects model. This standardisation was for visualisation of the data only. The plot demonstrates that the CUE decreases up to the CUE break-point temperature with a consistent temperature dependence, equating to an average activation energy (*E_a_^CUE^*) of -0.12eV. **(B)** Boxplot of the species-specific *E_a_^CUE^* values derived from the linear mixed effects model. The bold horizontal line corresponds to the median value, the top and bottom of the box correspond to the 75^th^ and 25^th^ percentiles and the whiskers extend to the largest and smallest values no greater or less than *1.5 ×* the interquartile range, beyond which the points are plotted as outliers.

It is important to note that our experiments were conducted under nutrient replete conditions. A recent study has suggested that the temperature sensitivities of photosynthesis and respiration (25) in some marine phytoplankton may decline under nutrient limitation and that the differential temperature sensitivities of photosynthesis and respiration may be negligible under limited conditions. This work however quantified the temperature sensitivities of photosynthesis and respiration at only 3 or 4 temperatures leading to estimates of thermal sensitivities with large error margins and a high probability of generating type II errors (i.e. accepting the null hypothesis of no difference in the thermal sensitivity of photosynthesis and respiration). Furthermore, measurements were made only under resource limited conditions precluding a quantitative comparison with nutrient replete conditions via the same methodology. Whilst we expect that the absolute values of the thermal sensitivities of photosynthesis and respiration are likely to decline under resource limitation, it is highly improbable that the intrinsic differences between photosynthesis and respiration documented in this study under nutrient replete conditions will be erased under nutrient limitation. Indeed, our analyses demonstrate that light limitation had a negligible impact on the temperature sensitivity of photosynthesis and in particular, the fundamental differences in the impacts of temperature on photosynthesis and respiration were preserved under light limited conditions (see Fig. 2 and Table 6, SI). We therefore anticipate that the supra-optimal declines in growth linked to temperature-driven decoupling between photosynthesis, respiration and biomass synthesis that we have shown here, apply equally under nutrient replete and limited conditions. Whilst large areas of the global ocean are under nutrient limited conditions for long periods (26), understanding the impacts of temperature under nutrient replete conditions (as we have done here) remains critically important because a large proportion of marine primary productivity occurs during episodic bloom events driven by short periods of increased nutrient concentrations (27–29). Clearly, significant further work is required to understand the interplay between temperature and nutrient availability on phytoplankton physiology and to assess whether the patterns we have shown here apply to conditions of nutrient limitation, given that current experimental evidence (25) is not sufficient to draw meaningful conclusions.

## Conclusions

Overall, our findings highlight marked similarities in the temperature dependence of photosynthesis and respiration across diverse taxonomic groups, spanning the cyanobacteria and red and green super families and suggest that common physiological trade-offs underpin the thermal tolerance of marine phytoplankton. We found that rates of respiration were more sensitive to temperature, had higher thermal optima and declined less abruptly past the optimum than those of photosynthesis. Consequently, the fraction of photosynthetic energy available for allocation to growth (the CUE) exhibited an accelerated decline with rising temperatures in a manner that was highly conserved among the 18 species investigated. We also found that the optimal temperature for growth coincided with, or was lower than, an inflection point in the temperature dependence of the CUE, which marked a transition that led to accelerated declines at high temperatures. These patterns suggest that universal metabolic constraints driven by the differential temperature sensitivity of photosynthesis and respiration play a key role in setting the limits of thermal tolerance of diverse marine phytoplankton. Our results therefore help pave the way for improving representations of phytoplankton biodiversity in models of ocean biogeochemistry by providing a process-based understanding of the factors that shape the limits of temperature tolerance for diverse species of marine phytoplankton, which can be used to aid predictions of immigration and local extinctions driven by global warming.

## Methods

### Culturing of marine phytoplankton strains

18 marine phytoplankton strains were obtained from CCAP (The Culture Collection of Algae and Protozoa) and RCC (Roscoff Culture Collection) between autumn 2015 and spring 2016. Strains of eukaryotic phytoplankton were selected from phylogenetic groups of both the red and green superfamilies (10, 11), in addition to two strains of cyanobacteria. We tried to work with organisms that had been well studied in the literature, were known to be globally abundant and play crucial roles for marine ecology and global carbon cycling. The strains were originally isolated from a range of latitudes and some have been in culture for up to 65 years (Table 1, SI). Stocks of each of the strains were cultured on their previous culture collection medium (Table 1, SI) using artificial sea water. The following media were used: Guillard’s F/2 and F/2 + Si, Keller’s K, K + Si and K/2, and PCR-S11 Red Sea medium (with Red Sea salts). All stock cultures were incubated in Infors HT incubators at 20°C, under a 12:12 hour light-dark cycle with a PAR intensity of 45-50 μmol m^2^ s^−1^ and shaken at 65RPM. Where possible we tried to obtain strains from the culture collections that matched, or were close to, these conditions. The red alga *Porphyridium purpureum* was an exception, which we cultured at 20-25 μmol m^2^ s^−1^. Cultures were kept under exponential, nutrient replete, growth conditions for ~ 2 months before any physiological data was collected.

### Measuring the thermal tolerance curve

For each species, a minimum of 3 technical replicates were inoculated with the same starting density into fresh growth medium across a range of temperatures (15°C - 37°C). Cell counts were made daily using flow cytometry (Accuri C6 flow cytometer, BD Scientific), and population density was tracked until cultures reached carrying capacity. Per capita growth rates (*μ*) were quantified from a modified Baranyi growth model without the lag phase(30), using non-linear least squares regression via the ‘nlsMicrobio’ package in R statistical software (v3.3.1). Models were fitted using the ‘nlsLoop’ function in the R github package ‘nlsLoop’. This draws on the ‘nlsLM’ function in the ‘minpack.lm’ R package, which uses a modified Levenberg-Marquardt optimisation algorithm. Model parameters were determined by running 2000 random combinations of estimated starting parameters, which were then selected using the Akaike Information Criterion (AIC) to determine the set of parameters that best characterised the data. Growth rates derived for each technical replicate at each growth temperature were then used to determine the thermal tolerance curves (Fig. 1A).

### Estimates of Cell Carbon and Nitrogen

For each species, an exponentially growing culture from the 20°C stock was divided into 3 technical replicates and centrifuged at 3500RPM, at 4°C for 30 minutes. The resultant pellets were rinsed with deionised water and re-spun 3 times to remove any artificial sea water residue. For the calcifying organisms (*Emiliania huxleyi, Gephyrocapsa oceanica, Thoracosphaera heimii* i.e. those with a calcium carbonate coccoliths) it was necessary to dissolve the extra-cellular inorganic carbon (31, 32). This was achieved by treating these pellets with 0.5 mL of 3M HCl for 1 hour before being rinsed with deionised water and re-pelleted. All pellets were freeze-dried using a CoolSafe (95-15 PRO, ScanVac) over 24 hours and then weighed to obtain dry weight. Samples were placed in tin cups and sent to Elemtex (Elemtex Ltd, Cornwall, UK, PL17 8QS) for elemental analysis of %C and %N using a SerCon Isotope Ratio Mass Spectrometer (CF-IRMS) system (continuous flow mode). For each technical replicate we then calculated the C:N ratio as well as μg C cell^−1^ (Table 3, SI).

### Measuring the metabolic thermal response curves

Measurements of photosynthesis and dark respiration were collected across a range of assay temperatures (7°C to 49°C) for a minimum of 3 biological replicates per species. We used a clark-type oxygen electrode as part of a Chlorolab 2 system (Hansatech Ltd, King’s Lynn, UK) to measure net rates of oxygen evolution in the light (net primary production, NP) and oxygen consumption in the dark (dark respiration); both in units of μmol O2 mL^−1^ s^−1^. All biological replicates were sampled from the stock cultures, which had all been growing at 20°C and were taken at the mid-logarithmic growth phase to ensure that the samples were not substrate limited. To improve the signal to noise ratio when measuring rates, all biological replicate samples were concentrated by centrifugation at 1500rpm, 20°C, for 15 minutes and re-suspended into an adequate volume of fresh growth medium. Prior to running a sample at each assay temperature, all samples were given ~ 15 minutes to pre-acclimate to the assay temperature in the dark before any data was collected. This also gave the electrode system sufficient time to stabilise before metabolic rates were measured. This was necessary for two reasons, i) as the sample adjusts to the assay temperature this will naturally cause changes in the dissolved oxygen concentration, ii) the electrode system results in oxygen signal drift, and this too is temperature dependent. We measured rates of oxygen depletion from 21 sterilised artificial seawater samples across a range of temperatures 4°C - 44°C and found that the impact of drift was minimised after ~15 minutes of stabilisation time. Nevertheless, signal drift was linearly temperature dependent after this time. To account for drift in our dataset we corrected all our raw data using the following empirically derived relationship:

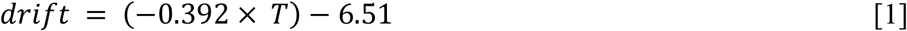

Where *T* is assay temperature (°C), and *drift* is the non-biological depletion in oxygen concentration measured in units μmolO_2_ mL^−1^ s^−1^ after approximately 15 minutes of stabilisation. The raw O_2_ flux data was then corrected by subtracting the estimated drift. Rates of net photosynthesis, measured as O2 evolution, were collected across a range of light intensities from 0 to 1800 μmol m^2^ s^−1^ with increments of 50 μmol m^2^ s^−1^ between 0 to 200 μmol m^2^ s^−1^, 100 μmol m^2^ s^−1^ between 200 and 1000 μmol m^2^ s^−1^, followed by 1200 μmol m^2^ s^−1^, 1500 μmol m^2^ s^−1^ and finally 1800 μmol m^2^ s^−1^. This enabled us to model a photosynthesis-irradiance (PI) curve for each assay temperature, and therefore obtain an estimate of light saturated net photosynthesis, *NP_max_*, see Eq. 2. Respiration (R) was measured as oxygen consumption in the dark, over a 3-minute period directly following the light response outlined above. The photosynthesis-irradiance curve was then quantified by fitting Eiler’s photoinhibiton model to the data using non-linear least squares regression (as described above) (33, 34):

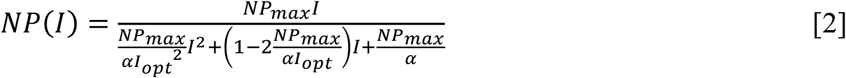

Where *NP*(*I*) is the rate of net primary production at light intensity, *I*, *NP_max_* is the maximum rate of *NP* at the optimal light intensity, *I*_opt_, and *a* is the rate in which *NP*increases up to *NP*_max_.

Light saturated gross primary production (*P*) was then calculated for each assay temperature as:

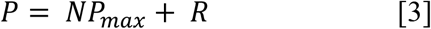

To investigate the effect of light limitation on the temperature dependence of photosynthesis we used Eq. 2 to determine the predicted *NP* at half the light saturated irradiance (0.5 × *I_opt_*). Thus replacing in *NP_max_* in Eq. 3 with this prediction we derived *P*_0.5_, a light limited value of gross primary production at half the saturating irradiance for each assay temperature response.

Metabolic rates were then converted from units μmol O_2_ mL^−1^ s^−1^ to μg C μg C^−1^ hour^−1^. We achieved this using the following equation:

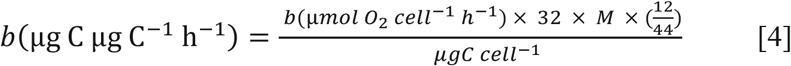

Where *b* is the metabolic rate (either *P* or *R*), 32 is the molecular weight of O_2_, *M* is a species specific assimilation quotient for CO_2_:O_2_ (35) which is used to describe consumption or fixation of C in the cell per unit of O_2_, and 12/44 is the ratio of molecular weight of C to CO_2_, thus 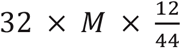 converts from *μmol O*_2_ to μ*gC*. Samples from each strain were analysed to determine species-specific μg C cell^−1^ values and the number of cells mL^−1^ was measured for each biological replicate using flow cytometry. The calculation of *M* is based on the assumption that NO_3_^−^ is the main nitrogen source in the growth medium and that there is a balanced growth equation, where:

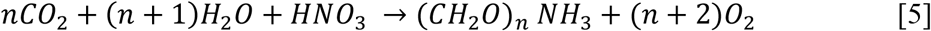

If the *C:N* ratio (*n*) of the phytoplankton is calculated in moles then the ratio of CO_2_:O_2_, or *M*, will be equal to *n/n*+*2* (35). Our calculated values of *M* ranged from ~0.71 to ~0.89 (Table 3, SI).

### Quantifying the thermal response curves

The thermal response curves for rates of growth, photosynthesis (at both saturated and half saturated irradiance) and respiration were quantified using a modified version of the Sharpe-Schoolfield equation (12, 13):

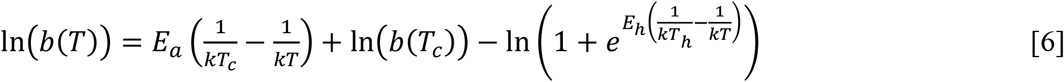

where *b* is either the rate of growth (d^−1^), photosynthesis or respiration (μg *C*μg C^−1^ h^−1^), *k* is Boltzmann’s constant (8.62×10^−5^ eV K^−1^), *E_a_* is the activation energy (eV), indicative of the steepness of the slope leading up to the thermal optima, *T* is temperature in Kelvin (K), *E_h_* is the deactivation energy which characterizes temperature-induced decrease in rates above *T_h_* where half the enzymes have become non-functional and *b*(*T_c_*) is rate normalized to an arbitrary reference temperature, here *T_c_* = 20°C (+ 273.15), where no low or high temperature inactivation is experienced. Eq. 6 can be used to derive an optimum temperature where the maximum rate is predicted:

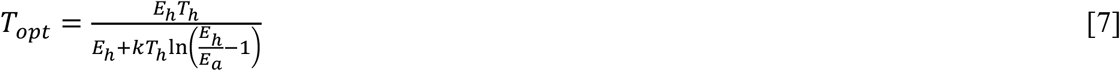

The parameters *b*(*T_c_*), *E_a_*, *E_h_*, *T_h_*, and *T_opt_*, can be considered as traits that characterise the unimodal response of biological rates to temperature change. We expect these traits to differ across the diverse taxa analysed in this study, owing to their diverse evolutionary histories and ancestral temperature regimes (given that they have been isolated from different latitudes/oceans). To test this assumption, we fitted the data for growth, photosynthesis and respiration across all species to Eq. 6 using non-linear mixed effects modelling with the ‘nlme’ package in R. We used separate analyses to assess the thermal responses of growth, photosynthesis and respiration. All models included each of the parameters in Eq. 6 as fixed effects, which quantify the average value of the parameter across all species and replicates. For the analysis of the thermal response of growth rate, we included ‘species’ as a random effect on each parameter, which quantifies species-specific deviations from the average across all species (i.e. the fixed effect) that are assumed to be normally distributed with a mean of zero. For the analyses of photosynthesis and respiration, we included ‘replicate’ nested within ‘species’ to account for the fact that we measured a minimum 3 replicate thermal response curves for each species. Here the random effect quantifies species-specific deviations from the fixed effects as well as those attributable to variance among the replicates of each species.

Because the Sharpe-Schoolfield equation can only take non-zero and positive rate values, in instances where either no observed growth rate, or a negative growth rate were measured (typically the highest and lowest temperature) we set the rate to the minimum value measured for the species in order to fit the model.

### Quantifying the carbon-use efficiency and modelling the break-point temperature

The carbon-use efficiency (CUE) was calculated as:

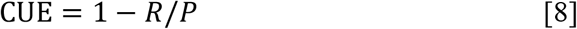

Due to the non-linear temperature response of the CUE, with accelerated declines at high-temperatures, we fitted a segmented linear regression model to estimate the break-point in the temperature response after which the CUE exhibited an accelerated decline. We fitted the segmented linear regression model to CUE values derived from the fitted Sharpe-Schoolfield curves for each species enabling us to derive an estimate of CUE at every 1°C increment across the range of assay temperatures where metabolic rates were measured for each species (Fig 2. Main text). We fitted the break-point model to the CUE values using the ‘segmented’ package in R, where the breakpoint estimate is defined in the segmented model as the intersection where there is significant difference in slopes (36), determined by the Davies test for performing hypothesis (37). It is for this reason that it was necessary to use the predicted values of respiration and photosynthesis to derive the break-point, as the measured data in most cases only provided one or two data points beyond the inflection point, and this would not have been sufficient to accurately model the second slope beyond this point (Fig. 3, Main text). The model returned an estimate of the CUE break-point temperature and the 95% confidence intervals surrounding this value for each species (Table 7, SI).

### Determining the temperature dependence of the CUE

We characterized the temperature dependence of the CUE up to the CUE breakpoint temperature for each species using the Arrhenius equation,

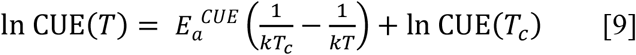

where In CUE(*T*) is the natural logarithm of the CUE at temperature *T* (in Kelvin), *E_a_^CUE^* is the apparent activation energy characterising the temperature dependence of CUE. We centred the temperature data using an arbitrary reference temperature *T_c_* = 283 K = 20°C, so that In CUE(*T_c_*) is the CUE at *T_c_*. We fitted Eq. 9 to all the measurements of CUE, up to the CUE break-point temperature identified for each species (Fig. 3 Main text, Table 7, SI) using a linear mixed effects model. This allowed us to derive an average value for *E_a_^CUE^* and In CUE(*T_c_*) across the 18 species. We also included random effects of ‘replicate’ nested within ‘species’ in the model to account for the fact we measured a minimum of 3 replicate responses of respiration and photosynthesis for each species. This allowed us to capture the species-specific and replicate specific estimates *E_a_^CUE^* and In CUE(*T_c_*).

## Acknowledgements

This study was supported by a grant from the Leverhulme Trust (RPG-2013-335) awarded to G.Y.-D., A.B. and N.S.

## Author contributions

S.B. and G.Y.-D conceived the study. S.B., G.Y.-D, A.B. and N.S. designed the experimental work. S.B., J.J. and C.-E.S. conducted the experimental work. S.B. and G.Y.-D analysed the data. S.B. and G.Y.-D wrote the manuscript and all authors contributed to revisions.

